# Fine-scale variation in microclimate across an urban landscape changes the capacity of *Aedes albopictus* to vector arbovirus

**DOI:** 10.1101/090613

**Authors:** C. C. Murdock, M. V. Evans, T. McClanahan, K. Miazgowicz, B. Tesla

**Affiliations:** Department of Infectious Diseases, College of Veterinary Medicine, University of Georgia, 501 D.W. Brooks Drive, Athens, GA 30602 U.S.A.; Odum School of Ecology, University of Georgia, 140 E. Green Street, Athens GA 30602 U.S.A.; Center for Tropical and Emerging Global Diseases, University of Georgia, 500 D.W. Brooks Drive, Athens GA 30602, U.S.A.; Center for the Ecology of Infectious Diseases, Odum School of Ecology, University of Georgia, 140 E. Green Street, Athens GA 30602, U.S.A.; Center for Vaccines and Immunology, College of Veterinary Medicine, University of Georgia, 501 D.W. Brooks Drive, Athens GA 30602, U.S.A.; University of Georgia Riverbasin Center, University of Georgia, 203 D.W. Brooks Drive, Athens, GA 30602, U.S.A.; Mathematics, University of Arkansas Little Rock, 2801 S. University Avenue, Little Rock AR 72204, U.S.A.

**Author notes:** **Corresponding author:** C. C. Murdock.

**Keywords:** Asian tiger mosquito, *Aedes*, vector, arbovirus, urban, climate, model, vectorial capacity

## Abstract

Most statistical and mechanistic models used to predict mosquito borne disease transmission incorporate climate drivers of disease transmission by utilizing environmental data collected at scales that are potentially coarser than what mosquito vectors actually experience. Temperature and relative humidity can vary greatly between indoor and outdoor environments, and can be influenced strongly by variation in landscape features. In the *Aedes albopictus* system, we conducted a proof-of-concept study in the vicinity of the University of Georgia to explore the effects of fine-scale microclimate variation on mosquito life history and vectorial capacity (VC). We placed *Ae. albopictus* larvae in artificial pots distributed across three replicate sites within three different land uses – urban, suburban, and rural, which were characterized by high, intermediate, and low proportions of impervious surfaces. Data loggers were placed into each larval environment and in nearby vegetation to record daily variation in water and ambient temperature and relative humidity. The number of adults emerging from each pot and their body size and sex were recorded daily. We found mosquito microclimate to significantly vary across the season as well as with land use. Urban sites were in general warmer and less humid than suburban and rural sites, translating into decreased larval survival, smaller body sizes, and lower per capita growth rates of mosquitoes on urban sites. Dengue transmission potential was predicted to be higher in the summer than the fall. Additionally, the effects of land use on dengue transmission potential varied by season. Warm summers resulted in a higher predicted VC on the cooler, rural sites, while warmer, urban sites had a higher predicted VC during the cooler fall season.

## 1. INTRODUCTION

Epidemics of dengue, chikungunya, and Zika are spreading explosively through the Americas creating a public health crisis that places an estimated 3.9 billion people living within 120 different countries at risk. This pattern began with the growing distribution of dengue virus (DENV) over the past 30 years, infecting an estimated 390 million people per year. More recent invaders, chikungunya (CHIKV) and now Zika virus (ZIKV), are rapidly following suit. CHIKV emerged in the Americas in 2013 and has caused 1.8 million suspected cases from 44 countries and territories (www.paho.org) to date. In 2015, outbreaks of Zika virus (ZIKV) have spread throughout the Americas, resulting in over 360,000 suspected cases, with likely many more going unreported.

Temperature is one of the key environmental drivers influencing the dynamics and distribution of these diseases^1–10^. Variation in temperature can profoundly impact mosquito population dynamics^11^, mosquito life history traits^12–18^ (such as survival, fecundity, duration of gonotrophic cycles, mosquito immune responses^19–22^), and measures of parasite / pathogen fitness (prevalence, titers, and the extrinsic incubation period)^1,10,23,24^. In addition to environmental temperature, variation in precipitation^25–27^ and relative humidity^28^ also drive vector-borne disease transmission.

Most statistical and mechanistic models used to predict mosquito borne disease transmission incorporate climate drivers of disease transmission by utilizing environmental data collected from general circulation weather models^1,29–32^, down-scaled weather data^33^, outdoor weather stations^34,35^, or remotely sensed land surface temperature data^36–38^. While working with these data is methodologically tractable, mosquitoes do not experience environmental variation at such coarse scales^39,40^. Temperature and relative humidity can vary greatly between indoor and outdoor environments^41,42^, and can be influenced strongly by variation in landscape features such as density of housing, housing material, vegetation cover, impervious surface cover, waste heat generation, and distance to water^18,28,43–48^. Thus, the microclimate a mosquito vector experiences will be dependent upon its dispersal ability (can be < 100 m for some species^49^) and the habitats it visits throughout life. In addition, many modeling efforts characterize environmental conditions through measures of mean monthly temperature, relative humidity, and precipitation. Yet, there is a growing body of theoretical and empirical work demonstrating that daily fluctuations in temperature, and likely relative humidity, are important for both mosquito and parasite / pathogen traits that mediate transmission ^1,2,5,43^. Thus, most studies characterizing environmental effects on vector-borne disease transmission are challenged by capturing the relevant metrics of environmental variation and the appropriate spatial and temporal scale.

We conducted a semi-field study examining differences in microclimate and mosquito life history traits across a heterogeneous urban landscape to address the above concerns. Specifically, 1) does mosquito relevant microclimate vary across an urban landscape, 2) how does this variation affect mosquito life history traits, and 3) what are the implications of microclimate variation for vectorial capacity? We investigated these questions in Athens-Clarke Country, GA, focusing on the invasive *Aedes albopictus* (Asian tiger mosquito) system due to its widespread distribution throughout the state^50^, as well as its role as an important vector for dengue, chikungunya, and Zika viruses in many parts of the world^51–54^.

## 2. METHODS

### 2.1 Site selection

We explored microclimate variation across three levels of land use categories characteristic of an urban landscape: urban, suburban and rural. We used an impervious surface map of Georgia generated by the Natural Resources Spatial Analysis Lab at the University of Georgia (http://narsal.uga.edu/glut/data-stats/georgia-impervious-surface-trends) for Athens-Clarke County, Georgia, U.S.A. to distinguish sites into these three land use categories. We defined urban, suburban, and rural sites as those with impervious surface scores within the following binned ranges: 55 – 100%, 5 – 50%, and 0%, respectively. We then created a new impervious surface map for Athens-Clarke County and selected three replicate sites within each land use category (Figure 1). Final site selection across Athens-Clarke County was ultimately constrained to sites that we could get permission to access. We did ensure that there was greater than 5 miles between sites, sites were interspersed across the county, and they were of the same area (30 m^2^, Figure 1).

**Figure 1.**
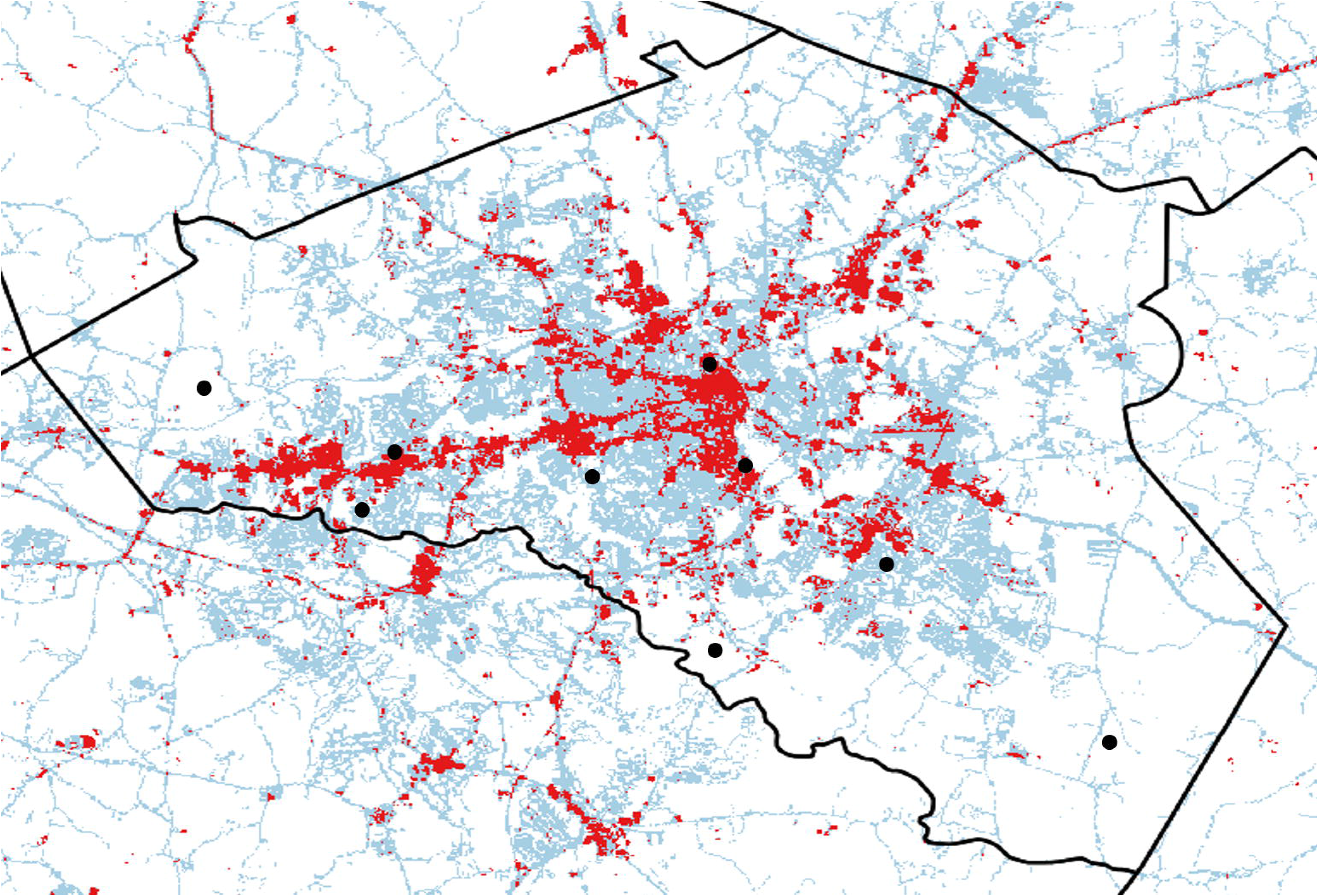
An impervious surface map of Athens-Clarke County, Georgia, U.S. modified from the Georgia impervious surface map produced by the Natural Resources Spatial Analysis Lab at the University of Georgia. Spatial pixels (30 m^2^) were binned according to proportion of impervious surface, with high, intermediate, and low proportion of impervious surface corresponding to urban (red), suburban (blue), and rural (white) sites, respectively. From this map, we selected three sites (black dots, 30 m^2^) from each land use class for the artificial pot experiments.

### 2.2 Larval development experiment

Within each site, we evenly distributed (10 m apart) and staked six black flower pots (Home Depot 480064-1001) in the ground at the base of vegetation (e.g. grass stands, brush, trees) in full shade. Within each pot, we placed a wide-mouth glass bell jar (∼1 L, Walmart, 550797441), and added 300 mL of leaf infusion and 30 first instar *Aedes albopictus* larvae. Leaf infusion was made a week prior to the start of the experiment. Live oak (*Quercus virginiana)* leaves were collected from the field and dried in an oven (50°C) for 72 hrs to ensure all water had evaporated from the leaf tissue. We then infused 80 grams of dried leaf material and 3 grams of a 1:1 yeast-albumin mixture in 20 L of deionized water for 3 days prior to use. To monitor variation in larval and adult mosquito microclimate across each site, we added a data logger (Monarch Instruments: RFID Temperature Track-It logger) to each jar and hung a logger (Monarch Instruments: RFID Temperature and Relative Humidity Track-It logger) in vegetation near each jar (∼3 feet above the ground). Loggers recorded instantaneous measurements of temperature and relative humidity every 10 min throughout the course of the study. Jars were then screened to prevent any emerging adult mosquitoes from escaping and a wire cage (8 in x 8 in) with plastic vinyl lining the roof was placed over top and staked into the ground to exclude animals and excess rainfall.

Pots were visited daily and any emerged adults were removed. We quantified the total number of adults emerging per day, and recorded the sex and wing length of each emerged adult. Wing length was used as a proxy of mosquito body condition because of its associations with female mosquito fecundity, survival, and vector competence for arboviruses^55–57^. One wing was taken from each individual upon emergence, mounted on a glass microscope slide using clear nail lacquer, and measured using a dissecting scope and micrometer eye piece. Measurements were taken from the tip of the wing (excluding fringe) to the distal end of the alula. This experiment was conducted twice, once in early summer (June 15-July 14, 2015) and once in the fall (September 7-October 10, 2015) to estimate any effects of season on our response variables.

### 2.3 Calculating per capita mosquito population growth rates (*r*)

We used the following equation (1) to calculate per capita intrinsic population growth rates (*r*) for each experimental pot across all sites^58^,

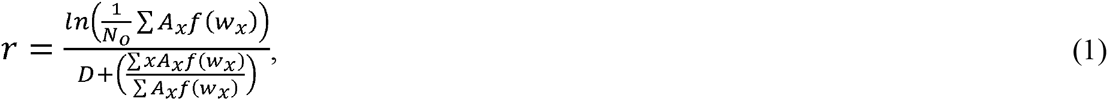
in which *N_o_* represents the initial number of females, *A_x_* the number of adult females emerging per day *x, w_x_* the mean wing length of females emerging on day *x, D* the delay between female emergence and first oviposition, and *f(w_x_)* predicts the numbers of female offspring produced by females of a given wing size. Because 1^st^ instar mosquito larvae cannot be reliably sexed, and 30 1^st^ instar larvae were deposited in each experimental pot, we assumed *N_o_* to be 15 females. We also assumed *D* = 14.2 days for *Ae. albopictus*^58^. We used the following linear function, *f*(*w_x_*) = −121.240 + 78.02*w_x_*, to describe the relationship between mean wing size and fecundity^59^.

### 2.4 Statistical analysis

To estimate the effects of microclimate and land use on the larval development and mosquito emergence rates, we used Cox proportional hazard models (R version 3.3.0, package ‘survival’) to assess how these predictors influenced probability of mosquito emergence across pots in each seasonal block (*summer* and *fall*). Each model included land use (*rural, suburban*, and *urban*) and the following microclimate covariates (*daily temperature mean, minimum*, and *maximum* in each experimental pot and average *daily relative humidity mean, minimum*, and *maximum*) as predictor variables. Additionally, to control for correlated observations, pot was included as cluster variables in the analysis. We achieved our final models by using a multidirectional stepwise selection method designed to minimize Akaike Information Criterion (AIC)^60^. All predictors included in final models were checked with the PH assumption. Influential observations and nonlinearity were investigated by removing one observation for each covariate and observing how much the regression coefficients changed and plotting the Martingale residuals against each covariate, respectively.

We used general linear mixed effects models (JMP^®^ Pro 12.1.0) to investigate the effects of seasonal block (*summer* and *fall*), land use class (*rural, suburban*, and *urban*), and the interaction on metrics of larval microclimate (average daily mean, minimum, and maximum temperature in each experimental pot and average daily mean, minimum, and maximum relative humidity), mosquito body size upon emergence (wing size), and the per capita mosquito population growth rate, *r*. Experimental pot was nested within site as a random factor within each model. Sex, and the interactions with seasonal block (*sex x block*) and land use (*sex x land use*), were also included as predictors in the model with wing size as the response variable. Model fit and assumptions of normality were assessed by plotting model residuals and quantile plots, and Tukey adjusted pairwise comparisons were run to compare differences across land use groups.

### 2.5 Estimating the effects of season and land use on transmission potential

To estimate how variation in relevant microclimate across different land uses and season might influence the ability of *Ae. albopictus* to transmit arboviruses, we used a dengue-specific vectorial capacity framework. Vectorial capacity (*VC*) is a mathematical expression (2) that integrates mosquito and pathogen life history traits that are relevant for transmission and describes the daily rate at which future infections arise from one infected human, given that all female mosquitoes feeding on that human become infected^10,61–63^:

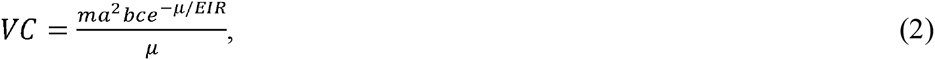
where *m* represents vector density, *a* is the daily probability of a human host being fed on by a vector, *EIR* is the extrinsic incubation rate of a pathogen, *μ* is the daily probability of adult mosquito mortality, *b*\**c* is vector competence, and *EIR* is the extrinsic incubation rate of a pathogen. The density of mosquitoes (*m*) was estimated with the following equation (3): 
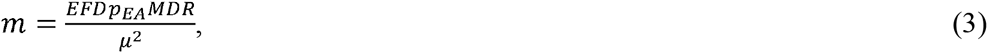
with *m* being comprised of the number of eggs laid per female per day (*EFD*), the egg to adult survival probability (*p_EA_*), the development rate of larvae (*MDR*), and adult daily probability of mortality (*μ*).

From our semi-field experiment, we can directly estimate the number of eggs laid per female per day (*EFD*) by taking the number of expected eggs laid per gonotrophic cycle based on body size, estimated by the linear relationship between eggs laid and wing length (y = 78.02x-121.24)^59^, divided by expected lifespan (1/*μ*) within each land use and season. From our survival analyses, we can estimate the probability of egg to adult survival (*p_EA_*) and the mosquito development rate (*MDR*). The *p_EA_* and *MDR* were estimated as the maximum percentage of adult females emerging across each site and the slope of the inflection point of the cumulative emergence curves, respectively. To estimate the parameters in vectorial capacity that we did not directly measure in our semi-field experiment (*a, b, c, EIR*, and *μ*) as a function of mean daily temperature (*T*) on our sites and across each season, we used either the Briere thermal equation (4) use to explain asymmetric, non-linear relationships,

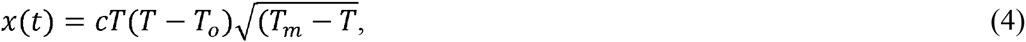
or the quadratic equation (5) used to explain symmetric relationships, 
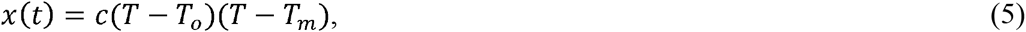
with *T_o_* as the daily minimum temperature, *T_m_* as the daily maximum temperature, and *c* as a fit parameter with values for these parameters taken from Mordecai et al. 2016 (under review)^35^. In order to estimate potential effects of variation in diurnal temperature ranges across our sites with land use and season on these parameters, we used rate summation^34^ (6) defined as 
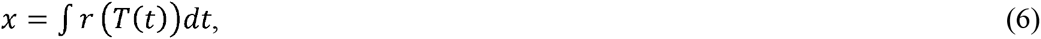
where a given trait (*x*) is defined as a rate (*r*) that adjusts instantaneously to temperature (*T*), which in turn is a function of time (*t*).

## 3. RESULTS

### 3.1 The effect of season and land use on mosquito microclimate

We found that the larval microclimate mosquitoes experienced significantly varied with time of season and with land use (Table 1, Figure 2). We did not observe any significant interactions between seasonal block and land use, suggesting that the effects of land use on mosquito microclimate were consistent across the summer and fall experiments. Due to larval data logger failure, we were unable to track daily water temperatures across a total of six pots (n=48 pots) in the summer and one pot (n=53) in the fall; however, as the failure was equally distributed across treatments, we do not believe this significantly affected our results.

**Figure 2.**
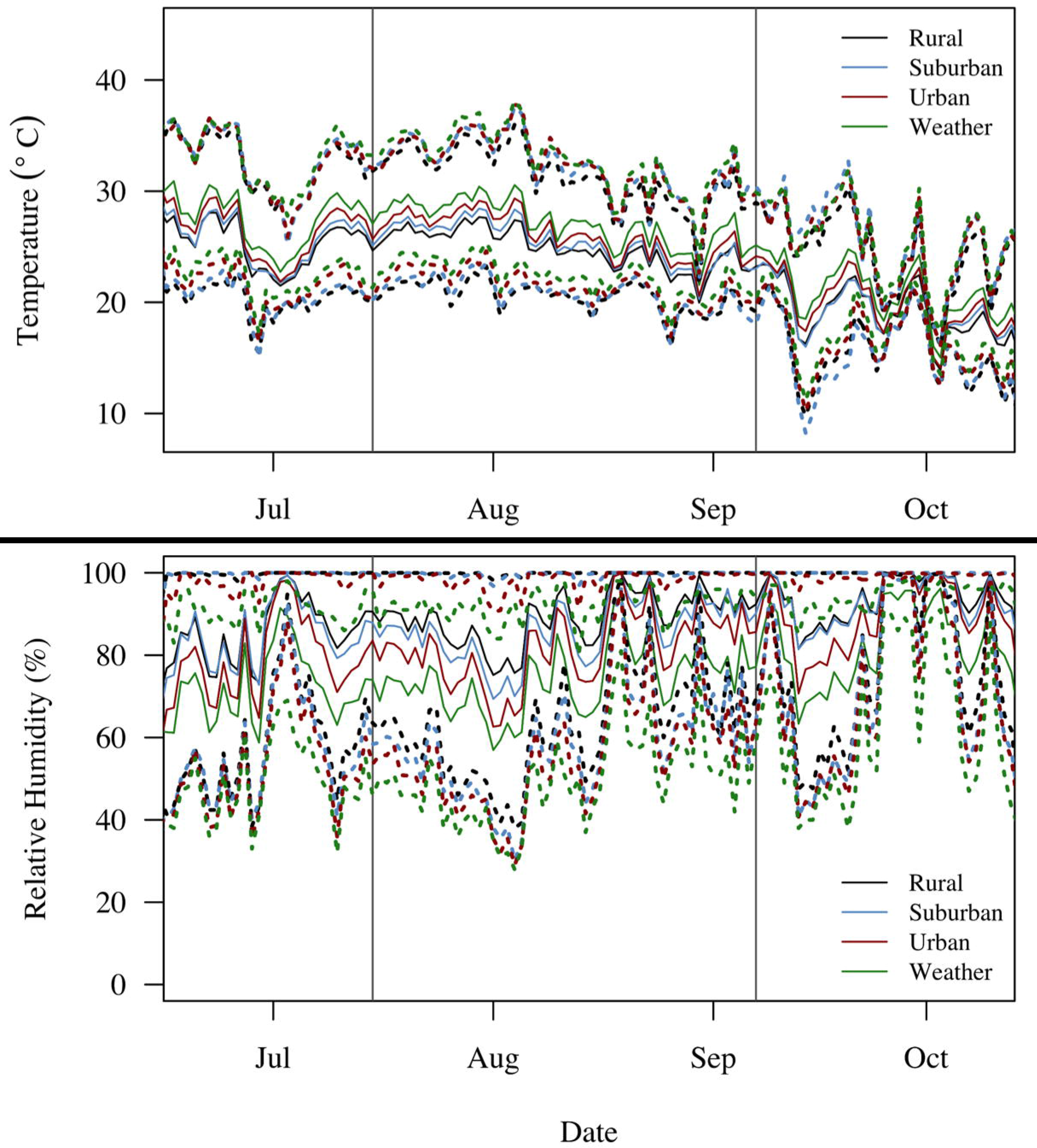
Ambient mean (solid lines), minimum (lower dotted lines), and maximum (upper dotted lines) daily temperature (**A**) and relative humidity (**B**) were recorded by data loggers across the duration of both experiments on urban (red), suburban (blue), and rural (black) sites and by the local weather station (green) on campus.

**Table 1.** Results from mixed effects models investigating the effects of season (summer vs. fall), land use (rural, suburban, urban), and the interaction on different daily measures of microclimate. Experimental pot nested within site was included as a random factor.

| Factors | *daily mean* |  |  | *daily minimum* |  |  | *daily maximum* |  |  | *diurnal range* |  |  |
| --- | --- | --- | --- | --- | --- | --- | --- | --- | --- | --- | --- | --- |
|  | d.f. | F | p | d.f. | F | p | d.f. | F | p | d.f. | F | p |
| *temperature* |  |  |  |  |  |  |  |  |  |  |  |  |
| season | 1 | 7329.58 | **<0.0001** | 1 | 6851.07 | **<0.0001** | 1 | 1215.54 | **<0.0001** | 1 | 69.91 | **<0.0001** |
| land use | 2 | 3.74 | **0.0307** | 2 | 4.62 | **0.0151** | 2 | 78.04 | **0.0016** | 2 | 5.40 | **0.0076** |
| season x land use | 2 | 1.05 | 0.3592 | 2 | 3.04 | 0.0585 | 2 | 13.20 | 0.1955 | 2 | 0.09 | 0.9174 |
| *relative humidity* |  |  |  |  |  |  |  |  |  |  |  |  |
| season | 1 | 774.17 | **<0.0001** | 1 | 594.33 | **<0.0001** | - | - | - | 1 | 564.93 | **<0.0001** |
| land use | 2 | 53.88 | **<0.0001** | 2 | 9.49 | **0.0003** | - | - | - | 2 | 4.70 | **0.0134** |
| season x land use | 2 | 2.68 | 0.0793 | 2 | 1.36 | 0.2726 | - | - | - | 2 | 2.49 | 0.0932 |

As expected, summer temperatures were on average higher than fall temperatures, with significantly higher daily mean (summer: 26.0°C; fall: 20.5°C) minimum (summer: 22.4°C; fall: 15.6°C), and maximum water temperatures (summer: 29.6°C; fall: 24.5°C). Additionally, experimental pots in the summer were subject to lower daily mean (summer: 82.8%; fall: 92.8%) and minimum relative humidity (summer: 55.9%; fall: 74.8%). We did not include maximum relative humidity in our analyses because the daily maximum relative humidity across all sites and seasons was close to 100%. These seasonal differences in daily temperature and relative humidity resulted in summer mosquitoes experiencing a lower diurnal temperature range (summer: 7.3°C; fall: 8.9°C) and higher diurnal relative humidity range (summer: 43.0%; fall: 25.0%) across all sites.

Urban sites were on average warmer than rural sites (Figure 2). Urban sites were characterized by higher daily mean temperatures (urban vs. rural, p = 0.0234; urban vs. suburban, N.S.; suburban vs. rural, N.S.) and maximum temperatures (urban vs. rural, p = 0.0011; suburban vs. urban, N.S.; suburban vs. rural, N.S.). Interestingly, daily minimum temperatures were similar across suburban and urban sites, with larvae on rural sites experiencing significantly lower daily minimum temperatures (rural vs. suburban, p = 0.0123; suburban vs. urban, N.S.; urban vs. rural, N.S.). Urban sites were also significantly drier. Urban sites had lower daily mean relative humidity (urban vs. suburban, p < 0.0001; urban vs. rural, p < 0.0001, rural vs. suburban, N.S.) and minimum relative humidity (urban vs. suburban, p = 0.0023; urban vs. rural, p = 0.0007). Finally, urban sites experienced on average wider fluctuations in diurnal temperature (urban: 8.5°C; suburban: 7.9°C; rural: 8.0°C) and relative humidity (urban: 36.1%; suburban: 33.2%; rural: 32.7%) ranges than suburban (p = 0.0088) and rural sites (p = 0.045).

While the daily climate data collected by the local weather station do track the daily variation in temperature and relative humidity recorded by data loggers (Figure 2), the local weather station did not accurately predict daily mean, minimum, maximum, and diurnal ranges of temperature and relative humidity across each land use (Figure 3). Further, the ability of the local weather station to predict urban, suburban, and rural microclimate varied qualitatively across seasons. For example, in the summer, local weather station data over predicted daily mean (by 1.3°C – 1.8°C), maximum, (by 3.0°C – 4.2°C) and diurnal temperature ranges (by 3.1°C – 3.7°C), while under predicting variation in the daily mean (by 6.8% to 13.3%), minimum (5.0% - 9.4%), and maximum relative humidity (6.4% - 8.2%) across all land uses (Figure 3). In contrast, in the fall, the local weather station was better able characterize daily mean (a difference of 0.3°C – 0.7°C), maximum (a difference of 0.8°C – 1.2°C), and the diurnal temperature range (-0.8°C to -0.4°C) across these sites. In the fall, like the summer, the local weather station continued to under predict the daily mean, minimum, and maximum relative humidity across urban, suburban, and rural sites. Interestingly, while the difference in relative humidity recorded by the local weather station and our data loggers was minimal in the summer (-1.3% - 1.2%), the local weather station in the fall marginally over estimates the relative diurnal humidity range (3.7% - 7.8%) in urban, suburban, and rural sites (Figure 3).

**Figure 3.**
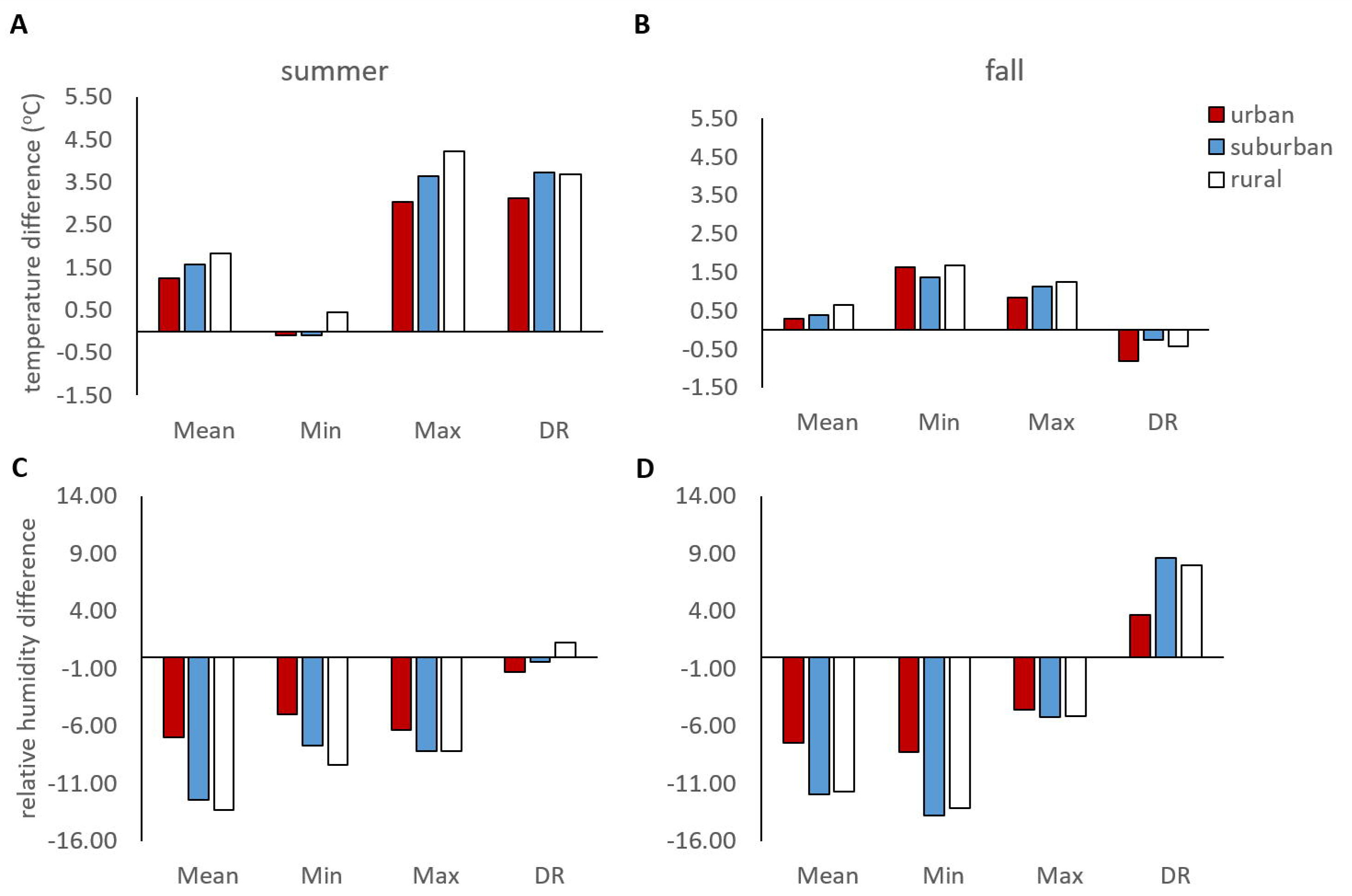
Differences between daily mean, minimum, and maximum values for temperature and relative humidity recorded by data loggers on urban (red), suburban (blue), and rural (white) sites in the summer (**A**, **B**) and fall (**C**, **D**).

### 3.2 The effect of microclimate, season, and land use on mosquito emergence

Overall, larval survival and the number of adult mosquitoes emerging were much higher in the fall than in the summer (Figure 4). Of approximately 1,620 first instar *Ae. albopictus* placed into the field during each experiment, we had a total of 318 females and 387 males successfully emerge during the summer replicate and 569 females and 623 males emerge during the fall replicate. Additionally, adults began to emerge at an earlier date in the summer (day 7) than in the fall (day 11). We found significant effects of land use on the likelihood of mosquito emergence in both the summer and fall, with a 44% and 47% decrease in the likelihood of mosquito emergence on urban sites relative to suburban and rural sites (which had similar likelihoods of mosquito emergence) in the summer, respectively (Table 2). There also was an effect of temperature and relative humidity on mosquito emergence in the summer and fall experiments, but interestingly these effects differed. Mosquitoes developing in the summer experienced an 18% *decrease* in the likelihood of emergence with each 1°C increase in the daily *minimum* temperature and a 7% decrease with each 1% increase in daily *mean* relative humidity (Table 2). In contrast, mosquitoes developing in the fall experienced a 28% *increase* in the likelihood of emergence with each 1°C increase in daily *maximum* temperature and a 19% decrease with every 1% increase in daily *maximum* humidity (Table 2). Together, these results suggest that higher temperatures on urban sites may decrease the likelihood of mosquito emergence through increased larval mortality, and that temperature variation throughout the day has qualitatively different effects on mosquito development and emergence in the summer than the fall.

**Figure 4.**
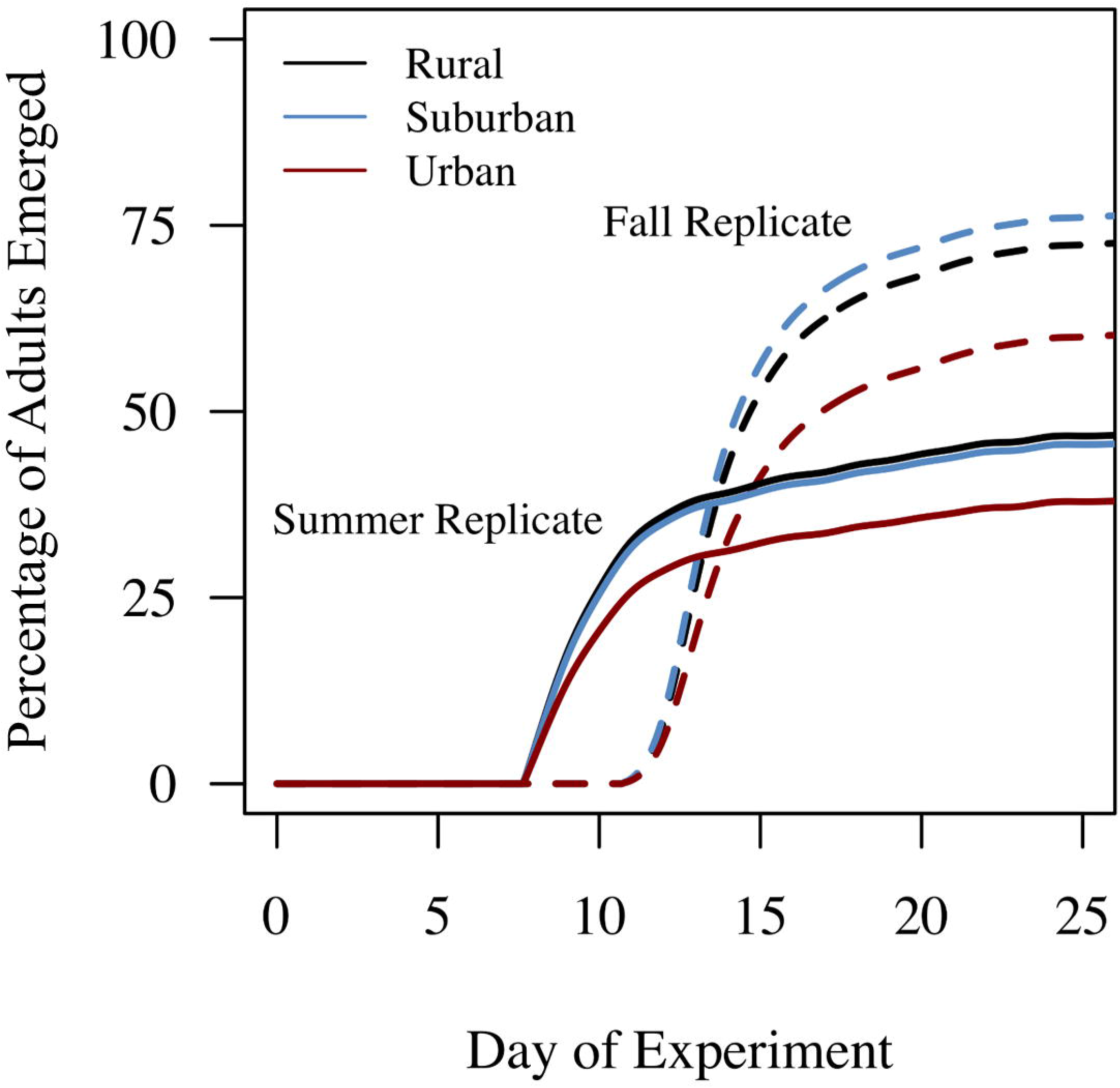
The cumulative percentage of mosquito adults emerging across urban (red), suburban (blue), and rural (black) sites in both the summer (solid lines) and fall (dashed lines).

**Table 2.** Final model results from a Cox proportional survival analysis estimating the effects of season (summer, fall), land use (rural, suburban, urban), and microclimate variables (daily mean, minimum, and maximum values of water temperature and ambient relative humidity) on the likelihood of mosquito emergence.

| Factors | *beta* | $e^{beta}$ | SE | z | p |
| --- | --- | --- | --- | --- | --- |
| *summer* |  |  |  |  |  |
| suburban | 0.0818 | 1.0852 | 0.1081 | 0.51 | 0.6095 |
| urban | -0.4206 | 0.6567 | 0.1527 | -2.15 | 0.0313 |
| daily minimum temperature | -0.1948 | 0.823 | 0.0872 | -2.13 | 0.0336 |
| daily mean relative humidity | -0.0693 | 0.9331 | 0.0212 | -2.64 | 0.0082 |
| daily minimum relative humidity | 0.0367 | 1.0374 | 0.0154 | 1.6 | 0.1093 |
| *fall* |  |  |  |  |  |
| suburban | 0.105 | 1.1107 | 0.7744 | 1.08 | 0.2804 |
| urban | -0.6299 | 0.5326 | 0.0972 | -2.77 | 0.0057 |
| daily average temperature | 0.1922 | 1.2119 | 0.1007 | 1.05 | 0.2932 |
| daily minimum temperature | -0.236 | 0.7898 | 0.0896 | -1.57 | 0.1173 |
| daily maximum temperature | 0.2518 | 1.2864 | 0.0769 | 2.25 | 0.0243 |
| daily maximum relative humidity | -0.2066 | 0.8134 | 0.0446 | -2.93 | 0.0034 |

### 3.3 Effects of microclimate, season, and land use on wing size and *r*

We found significant effects of sex, season, and land use on the size of emerging adult mosquitoes (Table 3, Figure 5). Overall, female mosquitoes were larger than male mosquitoes (females: 3.21 mm; males: 2.71 mm). Mosquitoes emerging in the summer were significantly smaller than those emerging in the fall (summer: 2.77 mm; fall: 3.15 mm), and mosquitoes developing on urban sites emerged as smaller adults (urban: 2.91 mm; suburban: 2.96 mm; rural: 3.01 mm) relative to rural sites (urban vs. rural, p = 0.0047; urban vs. suburban, N.S.; suburban vs. rural, N.S.). Interestingly, there were significant interactions between seasonal block and mosquito sex (*block x sex*) and land use (*block x land use*), suggesting the effects of season on mosquito body size differs for males and females and across land use. For example, female mosquitoes were always significantly larger than male mosquitoes, however this difference in body size was greater in the fall than the summer seasonal block. Further, there were no significant effects of land use on mosquito body size in the summer seasonal block, but mosquitoes emerging on urban sites were significantly smaller (urban: 3.09 mm; suburban: 3.14 mm; rural: 3.22 mm) than those on rural sites (urban vs. rural, p = 0.0003; urban vs. suburban, N.S.; suburban vs. rural, N.S.) during the fall.

**Figure 5.**
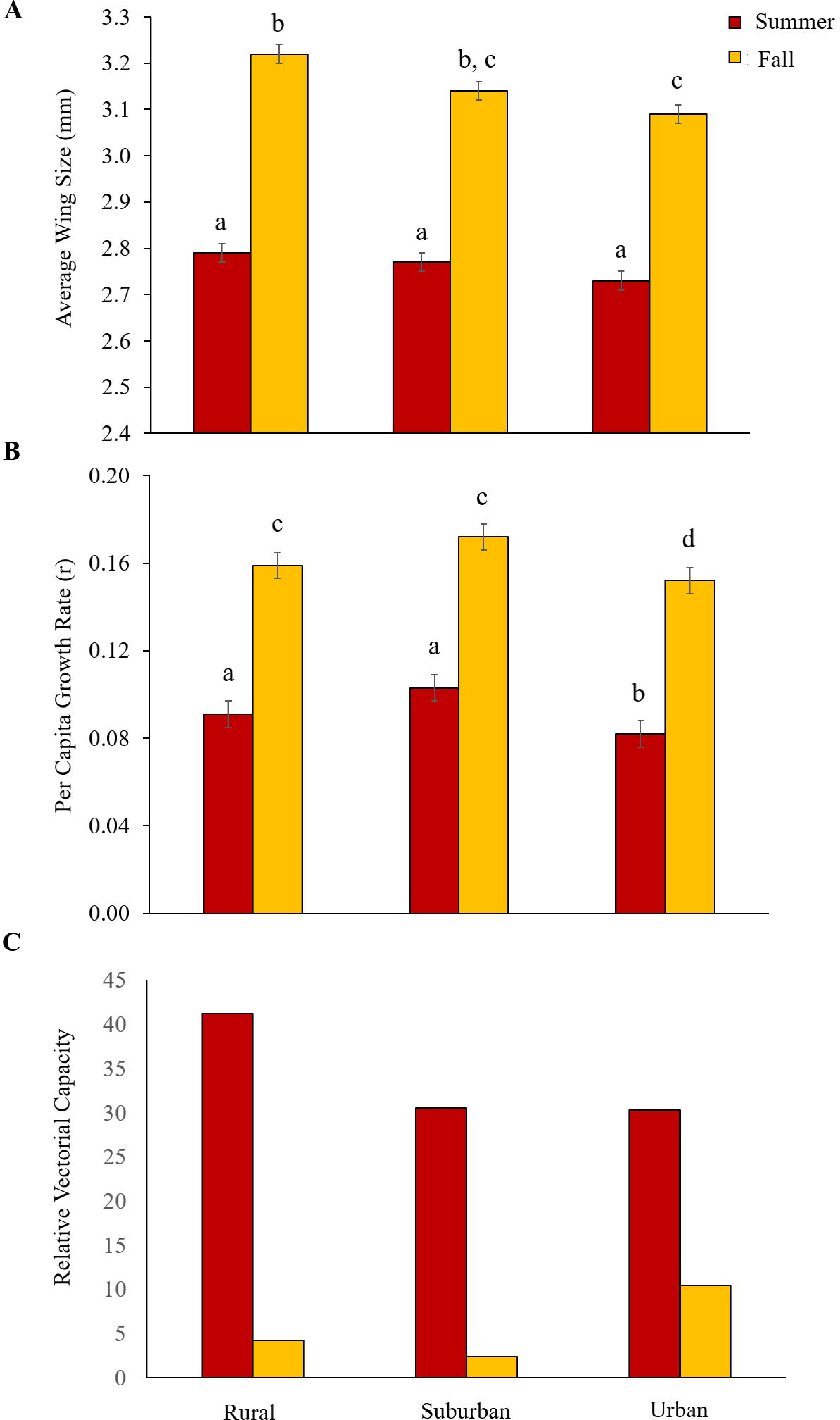
The effects of land use on mosquito body size (**A**), per capita mosquito growth rates (**B**), and relative vectorial capacity, or transmission potential (**C**) in the summer (orange bars) and fall (red bars). Error bars represent standard errors.

**Table 3.** Results from mixed effects model analysis with backward elimination investigating the effects of sex (male, female), season (summer, fall), land use (rural, suburban, urban) and all possible interactions on mosquito wing size, and the effects of season, land use and the interaction on mosquito per capita growth rates (*r*). Experimental pot nested within site was included as a random factor.

| Factors | *wing size* |  |  | *r* |  |  |
| --- | --- | --- | --- | --- | --- | --- |
|  | d.f. | F | p | d.f. | F | p |
| sex | 2 | 2670.26 | <0.0001 | - | - | - |
| season | 1 | 1590.73 | <0.0001 | 1 | 117.14 | <0.0001 |
| land use | 2 | 5.48 | 0.0069 | 2 | 3.58 | 0.0313 |
| season x land use | 1 | 4.52 | 0.011 | 1 | 0.50 | 0.6077 |
| sex x land use | 1 | 183.42 | <0.0001 | - | - | - |

Integrating the daily emergence and wing size data into equation (1), we identified significant effects of seasonal block and land use on mosquito per capita population growth rates (*r*, Table 4, Figure 5). Overall, the mosquito per capita growth rate was approximately two times higher in the fall (0.157) than the summer (0.090). Further, the mosquito per capita growth rate was on average significantly lower on urban sites (urban vs. suburban; 0.0269; urban vs. rural, N.S.; suburban vs. rural, N.S.).

### 3.4 The effect of land use and season on arbovirus transmission potential

We found mosquito transmission potential to vector dengue to vary across the summer season and with land use (Figure 5). Transmission potential was higher overall in the summer relative to the fall season. Interestingly, the effects of land use on mosquito transmission potential varied depending on time of season. The model predicts that during the hot summer, *Ae. albopictus* on rural sites have the highest transmission potential relative to suburban and urban sites. In contrast, in the cooler fall, mosquitoes on urban sites were predicted to have the highest transmission potential (Figure 5). Together these results demonstrate fine-scale variation in transmission potential can occur across an urban landscape, and seasonal shifts in microclimate can result in qualitatively different patterns of arbovirus transmission potential with land use.

## 4. DISCUSSION

To date, the majority of studies investigating the effects of urbanization on mosquito population dynamics and disease transmission have been sampling or modeling studies investigating how the distribution and abundance, feeding preferences, and incidence of diseases vectored by different mosquito species vary across land uses^46,64–73^. In contrast, there have been a handful of experimental studies in the field that mechanistically link observed variation in mosquito traits and metrics of disease transmission to sources of microclimate variation that exist across human-modified landscapes (*Anopheles spp.*^18,47,74^, *Culex pipiens*^45^, *Aedes albopictus*^75^). Our study, in combination with the previous work, demonstrates that relevant microclimate variation in the field (rather than coarser environmental manipulations in the lab) can translate into significant heterogeneity in mosquito life history traits, and ultimately disease transmission potential.

Across both the summer and fall, we observed urban microclimates to be significantly warmer and less humid than non-urban sites, which is reflective of the urban heat island (UHI) effect^76^. This is consistent with other studies showing that urban centers can have different temperature ^77–79^ and precipitation regimes ^80–82^ than surrounding areas. Urbanization results in significant modifications to the land-surface structure^44^, such as an increase in built surfaces and use of impervious materials (that absorb solar radiation), three dimensionality of the landscape (minimizing air flow), and decreases in vegetative cover (decreased shading and evaporative cooling). Additionally, urban centers produce more waste heat^44^. These shifts in temperature associated with UHIs are also associated with changes in other weather variables, such as precipitation regimes, wind speed, relative humidity, etc. all of which can impact the heat budget of organisms living in urbanized landscapes ^83^. Urban heat island effects in other systems have led to shifts in organism phenology (plants)^84–86^, life history (e.g. insect pests, ants, fruit bats)^87–90^, and overwintering behavior (mosquitoes)^79^, all of which can have significant implications for vector-borne disease transmission^72,79^. Further, because our study site (Athens, Georgia) is a relatively small city, the observed effects of land use on fine-scale variation in microclimate were subtle, suggesting that these effects could in fact be much larger in bigger cities. Larger cities in the United States, for example, can have mean temperatures that are on average 3°C higher than non-urban areas (with the exception of drier areas)^76^, and urban cores in mega-cities like New Delhi, India can differ by 10°C relative to surrounding non-urban areas^79^.

Despite the subtle effects of land use on mosquito microclimate, we still observed noticeable effects on larval survival, adult mosquito body sizes, intrinsic population growth rates, and overall transmission potential. This reinforces findings from a diversity of laboratory studies on *Ae. aegypti* and *Ae. albopictus* demonstrating the effects of relatively large changes in mean temperature^1,13,15,24,91–98^ and diurnal temperature range^1,7,99–101^ on a diversity of mosquito life history traits (e.g. survival, biting rate, fecundity, larval development, vector competence, and viral extrinsic incubation period). We found mosquitoes developing on urban sites experienced lower survival in the larval environment (approximately a 45% decrease in the probability of emergence across both seasons) and emerged as smaller adults than on non-urban sites, which could be due to urban sites being in general warmer than non-urban sites. Other similar studies report increases in mosquito development times^45,75^ on urban sites and an increase in adult mosquito emergence^75^, which we did not observe. The decrease in probability of emergence combined with smaller adult body size resulted in slightly lower per capita growth rates in the summer and fall on urban sites.

Surprisingly, different components dictating the diurnal range of temperature and relative humidity were important for larval survival. Overall, in the hot summer, the probability of adult mosquito emergence decreased with higher daily thermal minimums. In contrast, in the cooler fall, increases in the daily maximum temperatures corresponded to increases in the number of adults emerging. Despite having higher average daily thermal maximum temperatures relative to non-urban sites, mosquitoes developing on urban sites still experienced higher larval mortality in the fall. This suggests other sources of variation with land use might also be important for larval survival (biotic activity in larval environments, exposure to insecticides, etc.)^102,103^ on these sites. Relative humidity was also important for the probability of adult emergence across these sites, and like temperature, different metrics of relative humidity were important across different seasons. Interestingly, in both the summer and fall, increases in either the daily relative humidity mean or maximum resulted in proportional decreases in the probability of adult emergence. To the best of our knowledge, this is the first report of variation in relative humidity affecting the likelihood of larval survival and adult emergence.

Variation in daily temperature and relative humidity, as well as the observed variation in mosquito body size with land use and season, could have significant implications for other, unmeasured mosquito traits that are important for arbovirus transmission. For example, variation in both mean temperature and diurnal temperature range in the lab have been shown to impact the daily probability of adult survival (*μ*), female gonotrophic cycles and biting rates (*a*), the number of eggs females produce per day (*EFD*), vector competence (*bc*) and the extrinsic incubation period (*EIP*) for a diversity of mosquito species and pathogens (e.g. *Anopheles, Culex, Aedes*)^1,2,7,10,12,43,100,101,104,105^. Modeling studies have linked increased precipitation and relative humidity to increased disease incidence (e.g. dengue and malaria)^106–110^, likely through the negative effects of low relative humidity (e.g. < 40% relative humidity) on mosquito longevity^111^ and activity^112^. However unlike environmental temperature, the mechanisms of how relative humidity effects mosquito life history in general and disease transmission have been less well explored. Finally, variation in mosquito body size with land use and season could further compound the effects of temperature and relative humidity variation on these traits. The decreased body size of mosquitoes on urban sites could have negative implications for the daily probability of adult survival (*μ*)^113–115^ and egg production^95,116,117^, which in turn could impact the number of eggs females produce per day (*EFD*)^59^.

To explore how the effects of variation in daily temperature and diurnal temperature range could impact arbovirus transmission, we used a temperature dependent vectorial capacity equation parameterized for *Ae. albopictus*^35^ to predict how dengue transmission potential varies across urban, suburban, and rural sites and with season. While the vectorial capacity formula ignores some potentially important sources of variation (e.g. underlying the mosquito-human interaction), it provides a framework for estimating the relative importance of key mosquito / pathogen parameters and the effects of environmental variation on these parameters^1,43,118^. Interestingly, relative vectorial capacity was lower in the fall relative to the summer despite the fact that per capita mosquito population growth rates were predicted to be higher in the fall due to increased mosquito survival in the larval environment and larger body sizes. This is likely due to the negative effect of cooler temperatures on mosquito biting rate, the extrinsic incubation period of dengue, and the probabilities of transmission^35^. We also found arbovirus transmission potential to vary with land use, and the effects of land use on vectorial capacity depended on time of season. These results suggest that the environmental suitability for arbovirus transmission will be dependent upon the shape of the non-linear relationships mosquito and pathogen traits share with temperature, the daily average habitat temperatures and their proximity to the thermal optimum of this non-linear response, and how the effects of daily temperature fluctuation integrate with daily mean habitat temperatures to impact trait performance, and ultimately transmission potential.

This study captures how mosquito life history, potential population growth rates, and transmission potential respond to variation in microclimate with land use and season. However, there are many other factors that could vary, in addition to microclimate, with land use that could impact these variables. For example, variation in quantity and quality of larval habitat, adult resting habitat, access to hosts, and insecticide application with land use will also likely influence mosquito population dynamics, densities, and transmission potential^69,119–122^. Additionally, while environmental conditions shape the potential distribution and magnitude of disease vectors, socio-economic and demographic factors (e.g. outdoor recreation, housing quality, etc.) determine the level of human exposure and the realized transmission risk^123,124^. For example, even though transmission potential is predicted to be lower in the fall than the summer, seasonal changes in human behavior may result in higher transmission risk in the fall, when cooler temperatures encourage more outdoor activity.

Most studies that consider the role of climate in vector-borne disease transmission use climate data reported from local weather stations. Our proof of concept study demonstrates that the climate conditions captured by local weather station data do not reflect the microclimates mosquitoes experience, and that subtle variation in mean and diurnal ranges of temperature and relative humidity can lead to appreciable variation in key mosquito / pathogen life history traits that are important for transmission. Greater effort is needed to quantify the activity space mosquitoes occupy and the conditions of relevant transmission environments. This will not only be important for predicting variation in transmission potential and risk across seasons, geographic regions, and land uses, but also for building in realistic environmental variation in future laboratory work on mosquito-pathogen interactions.

## 5. ACKNOWLEDGEMENTS

We thank Mark Brown and Anne Elliot for intellectual and facility support of this project by providing us with *Aedes albopictus* for our experiments. We also thank John Drake, members of the Murdock lab group, and Maria Huertas-Diaz for intellectual and personnel support of this project. This project was funded partially through the Population Biology of Infectious Diseases NSF REU program, as well as the Department of Infectious Diseases and the Odum School of Ecology, University of Georgia. Any opinions, findings, and conclusions or recommendations expressed in this material are those of the author(s) and do not necessarily reflect the views of the National Science Foundation

